# Deep Domain Adversarial Neural Network for the Deconvolution of Cell Type Mixtures in Tissue Proteome Profiling

**DOI:** 10.1101/2022.11.25.517895

**Authors:** Fang Wang, Fan Yang, Longkai Huang, Jiangning Song, Robin B. Gasser, Ruedi Aebersold, Guohua Wang, Jianhua yao

## Abstract

Cell type deconvolution is a computational method for the determination/resolution of cell type proportions from bulk sequencing data, frequently used for the analysis of divergent cell types in tumor tissue samples. However, deconvolution technology is still in its infancy for the analysis of cell types using proteomic data due to challenges with repeatability/reproducibility, variable reference standards and the lack of single-cell proteomic reference data. Here, we developed a novel deep learning-based deconvolution method (scpDeconv) tailored to proteomic data. scpDeconv uses an autoencoder to leverage the information from bulk proteomic data to improve the quality of single-cell proteomic data, and employs a domain adversarial architecture to bridge the single-cell and bulk data distributions and transfer labels from single-cell data to bulk data. Extensive experiments validated the performance of scpDeconv in the deconvolution of proteomic data produced from various species/sources and different proteomic technologies. This method should find broad applicability to areas including tumor microenvironment interpretation and clinical diagnosis/classification.

## Introduction

The quantitative partitioning of cell types and the estimation of their relative proportions in a tumor sample is of important clinical and prognostic significance. E.g. it can provide critical information for chemo- and immune-therapy^1–3^. Bulk sequencing technologies generally refer to lysing and sequencing all cells in a particular tissue, organ or organ system, and the subsequent measurement of the average abundance of genes/proteins. Bulk sequencing thus ignores the cellular heterogeneity of the sample^4^. The development of single-cell (transcriptome/proteome) sequencing technologies allows the isolation of single cells from tissue(s) for analysis and measurement of the respective molecules^5,6^. Despite the conceptual advantages of this technology, it has been challenging to apply it to explore cellular heterogeneity in tumor microenvironments in large sample cohorts due to the relatively high cost and incompatibility with samples that are formalin-fixed or paraffin-embedded tissue^7,8^. Therefore, computational methods have been developed to infer cell types and their proportions from transcript expression profiles from bulk tissue samples^7–10^, providing a relatively low-cost and convenient way of studying the cellular composition of tumor microenvironments in a clinical context.

The deconvolution of cell types for transcriptomic data derived from bulk tissue samples^4,8,11^ generally assumes that the mixture of cells can be modelled as a system of linear weight equations, therefore non-negative matrix factorization can be applied to estimate cell type proportions^8^. However, in reality, the expression profiles within each cell type are heterogeneous e.g. due to different cell cycle and biochemical states of tested cells, the microenvironment and background in sequence data. The resulting feature variability impedes high quality modelling using a linear distribution assumption. For this reason, alternative deep-learning-based methods consisting of the following steps have been proposed: i) collect scRNA-seq data with corresponding cell type labels; ii) train a deep learning model with labelled scRNA-seq data as reference; and iii) infer the corresponding cell type proportions of bulk data with the trained model^9,12,13^. In reality, these deep learning models were shown to be prone to overfitting on the reference datasets and failed on unseen bulk data from another source^9^. As a remedy, some methods were proposed to finetune the model on new datasets with known cell type proportions to improve performance^9^. However, the finetuning strategy limits the methods’ generalization and is impractical in real scenarios since clinical bulk data usually lack reliable cell type proportions beforehand.

To date existing deconvolution methods only focused on transcriptomics or genomics. No attention has been paid to deconvolving bulk proteomic data despite of an urgent need for algorithms that perform this task. The proteome provides critical biological and clinical information ranging from understanding biological mechanisms to discovering drug targets^14^. Furthermore, the concentration of most proteins cannot be accurately represented by the transcription/expression level of corresponding genes due to complex post-transcriptional regulation, RNA/protein degradation and post-translation modification^15–17^. Meanwhile, most proteins are present at over 1000-fold higher copy numbers than the corresponding transcripts and express much wider dynamic range of abundance than transcripts, and therefore show higher potential to distinguish cells and cell states^15,18^ and to provide unique biochemical insights. As a consequence, single cell proteomic data with annotated cell type labels are necessary for deconvolving bulk proteomic samples.

By far the most commonly used single-cell proteomic technologies (e.g., CyTOF, FACS) are mainly based on molecularly tagged antibodies which can detect a few to dozens of surface proteins or marker proteins in parallel in a high number of cells^22^. The limited number of proteins detected, however limits CyTOF data as a basis for cell deconvolution. With the advent of mass spectrometry-based single-cell proteomic technologies (e.g., SCoPE), the number of proteins that can be detected has significantly increased to up to 1000-3000 intracellular proteins and thus expanded both the number and types of proteins detected^19–21,23,24^. However, the processing of sufficient numbers of cells to generate data to support cell type deconvolution from bulk data remains challenging. Further, compared with transcriptomic data, single cell proteomic data to train and benchmark deconvolution algorithms face a number of additional challenges. These include significant background noise in single cell proteomic data, poor data quality and marked variation depending on analysis run and/or technology, and limited proteome coverage. Furthermore, the existing deconvolution methods were designed for transcriptomic data and are not directly applicable to proteomic data, because i) different distributions and value ranges between protein abundance and transcript expressions^15^. Specifically, most existing deconvolution methods for transcriptomic data are based on an assumed distribution, such as a Poisson or negative binomial distribution which are recognized as unsuitable for proteomic data^4,11^, ii) much lower coverage of proteome detected by single cell proteomics, compared with bulk proteomics^25,26^, a situation that is not considered by existing deconvolution methods, and iii) for single-cell proteomic data and bulk proteomic data,”batch effects” (variability among distributions) are pronounced, which is rarely considered by existing deconvolution methods^27^.

Here, to tackle these challenges, we propose a novel deep-learning-based method, scpDeconv, to estimate cell type proportions for tissue proteomic data, with single cell proteomic data from similar cohorts as reference. Based on the framework of mix-up strategy, autoencoder and domain-adversarial neural network, scpDeconv can circumvent the need to generate huge simulation datasets, make efficient use of single-cell proteomic data, impute the missing low-abundance proteins with autoencoder, and establish a deconvolution module by domain adversary training, thereby improving the model’s performance, particularly in proteomic data and real-world scenarios. The deconvolution approach developed will give new insights and potential to re-analyze clinical tissue proteomic data, reveal tumor microenvironment from the proteome level, and discover new clinical diagnostic markers.

## Results

### Overview of the deconvolution algorithm scpDeconv

In this study, we developed scpDeconv to infer cell proportions from bulk proteomic data with single cell proteomic data as reference. To fit for the characteristics of proteomic data, we blended mix-up strategy, autoencoder and domain adversary neural network to assemble the overall framework of scpDeconv (Figure 1).

**Figure 1.**
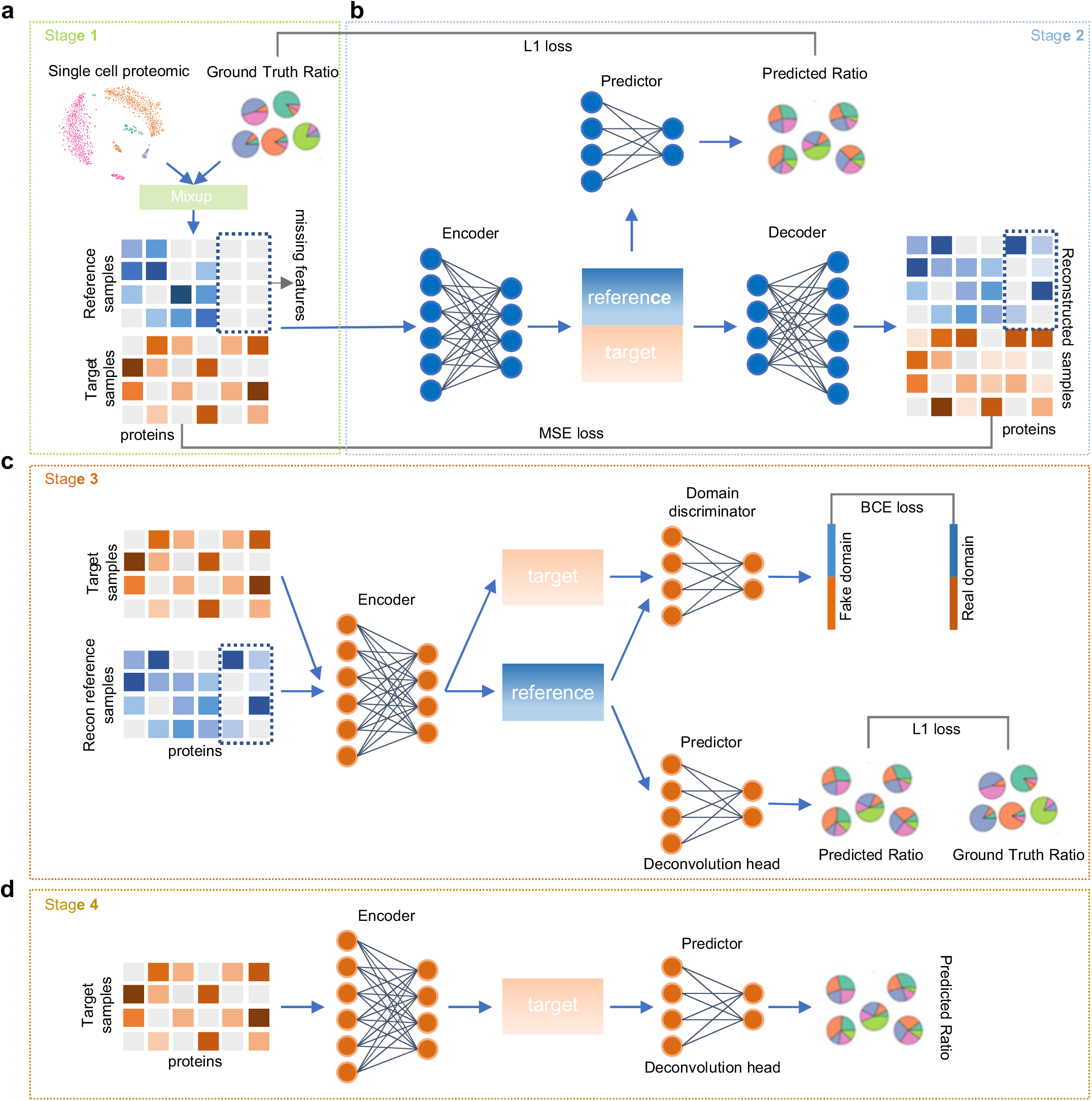
The architecture of our proposed method. a, Stage 1: reference data preparation. The mix-up process that combines the proteome profiles of different cell types by random proportions. b, Stage 2: missing features imputation and data quality improvement. The encoder-decoder scheme to close the gap between the reference data and the target data and impute the mixed-up single-cell proteomic data generated in Stage 1. Reconstruction loss (MSE loss) and deconvolution loss (L1 loss) are applied for model training. c, Stage 3: domain-adversarial training and deconvolution. The domain-adversary training architecture of neural network. The domain discriminator is used to discriminate whether the latent features are encoded from the reference data or from the target bulk proteomic data. The deconvolution head is used to predict the proportions of cell types. d, Stage 4: cell type proportion inference for target samples. Target bulk proteomic data are inputted to the trained encoder and deconvolution head from stage 3 to infer the proportions of cell types.

In stage 1 of scpDeconv (Figure 1a), we collected single cell proteomic datasets annotated with labelled cell types from the same species as the analyzed bulk tissue proteomic data (termed target data to be deconvolved). We then randomly mixed up single-cell proteomic data to construct pseudo-bulk data with corresponding cell type proportions as reference data.

In stage 2 of scpDeconv (Figure 1b), we applied an autoencoder model to impute missing low-abundance proteins in the reference data by borrowing information from target data. To be specific, this autoencoder consists of an encoder, a predictor and a decoder, uses both reference data and target data as input, and is trained with the reconstruction loss calculated between the raw input and the reconstructed output and the prediction loss calculated between the ground truth ratios and the predicted cell type proportions for reference data. In this way, by jointly training on labelled reference data and unlabelled target data, we can close the gap between reference data and target data, improve the quality of reference data, and impute low-abundance proteins which are undetected in single cell proteomic data.

In stage 3 of scpDeconv (Figure 1c), the imputed reference data and raw target data are sent to a shared encoder and the extracted features are passed through a domain discriminator for generative adversarial training, thereby facilitating the learning of domain-invariant features by the encoder. A deconvolution head is employed to predict cell type proportions from the extracted domain-invariant features. In this way, the method can efficiently improve the domain generalizability to overcome diverse distributions from single-cell proteomic data and bulk proteomic data.

Finally, in stage 4 of scpDeconv (Figure 1d), the corresponding cell type proportions of target proteomic samples are estimated by deep neural network trained in stage 3, and then utilized for re-analysis of biological mechanism or tumor microenvironment of specific tissue samples.

Overall, scpDeconv is designed for a real scenario in which bulk proteomic data from clinical or biological specimens are deconvolved to estimate the proportions of cell types constituting the tissue. This is accomplished by the use of prior single cell proteomic data from particular species and tissues and the annotated cell types as reference. It can be expected that with the present advances in the single cell proteomic technologies, a rapidly increasing number of high quality single cell proteomic data from various species and tissues will become available, thus increasing the reach of the presented method.

### Quantitative evaluation of cell type deconvolution performance

To quantitatively evaluate scpDeconv’s performance of cell type deconvolution, we carried out benchmarking calculations by simulating 3 different cases of increasing complexity. These are, i), cell type deconvolution across modalities using human breast atlas datasets consisting of scRNA-seq data and CyTOF data from the same source and involving 6 shared cell types within breast tissue; ii), cell type deconvolution across different data sources using 2 murine cell line datasets with 3 shared cell types generated with different single cell proteomic technologies, and iii), cell type deconvolution using integrated datasets from 5 human cell line datasets acquired by different single cell proteomic technologies from 4 shared cell types. In each case, we carried out the following steps to assess the deconvolution performance. First, we chose one or more single cell proteomic datasets to construct pseudo-bulk proteomic data with known cell type composition as reference data. Second, we also used similar single cell proteomic data in the same case to simulate pseudo-bulk proteomic data with true cell proportions as target data. Third, we employed scpDeconv to predict cell proportions of these target data. Finally, we compared scpDecov with four representative and state-of-the-art deconvolution methods developed for transcriptomic data, involving the deep-learning-based method (Scaden)^28^, the linear-regression-based method (MuSiC)^4^, the Bayes-based method (BayesPrism)^29^ and the distribution-hypothesis-based method (DestVI)^30^.

As shown in Figure 2a and Extended Data Figure 1a, evaluating by Lin’s concordance correlation coefficient (CCC), Root Mean Square Error (RMSE), and Pearson product-moment correlation coefficient (*r*), scpDeconv significantly outperformed Scaden, MuSiC, BayesPrism and DestVI in all the simulated cell type deconvolution cases. This is mainly due to the rational design of imputation to overcome the problem of missing low-abundance proteins in reference single-cell proteomic data, and the domain adversary training tailored for distribution differentiation between reference data and target data. The performance comparisons of the different tools provided further insights, as follows: i), compared to cases where cell type distribution data from the same source were available, exemplified the human breast atlas data, the performance of Scaden dropped when the target data were derived from samples with unseen cell distributions as was the case with murine cell line data from two different sources. This feature limits the general applicability of Scaden; ii), The performance of MuSiC was hampered by its heavy dependency on the consistency of gene weights across subjects, a feature that might be highly inconsistent between reference and target data; iii), BayesPrism showed poor performance due to its reliance on parameter estimation which required high-quality training data that are challenging to generate with present single cell proteomic technologies; vi), due to rigorous assumption about the distribution of transcripts in transcriptomic data, DestVI showed the worst performance and was hard to migrate to proteomics.

**Figure 2.**
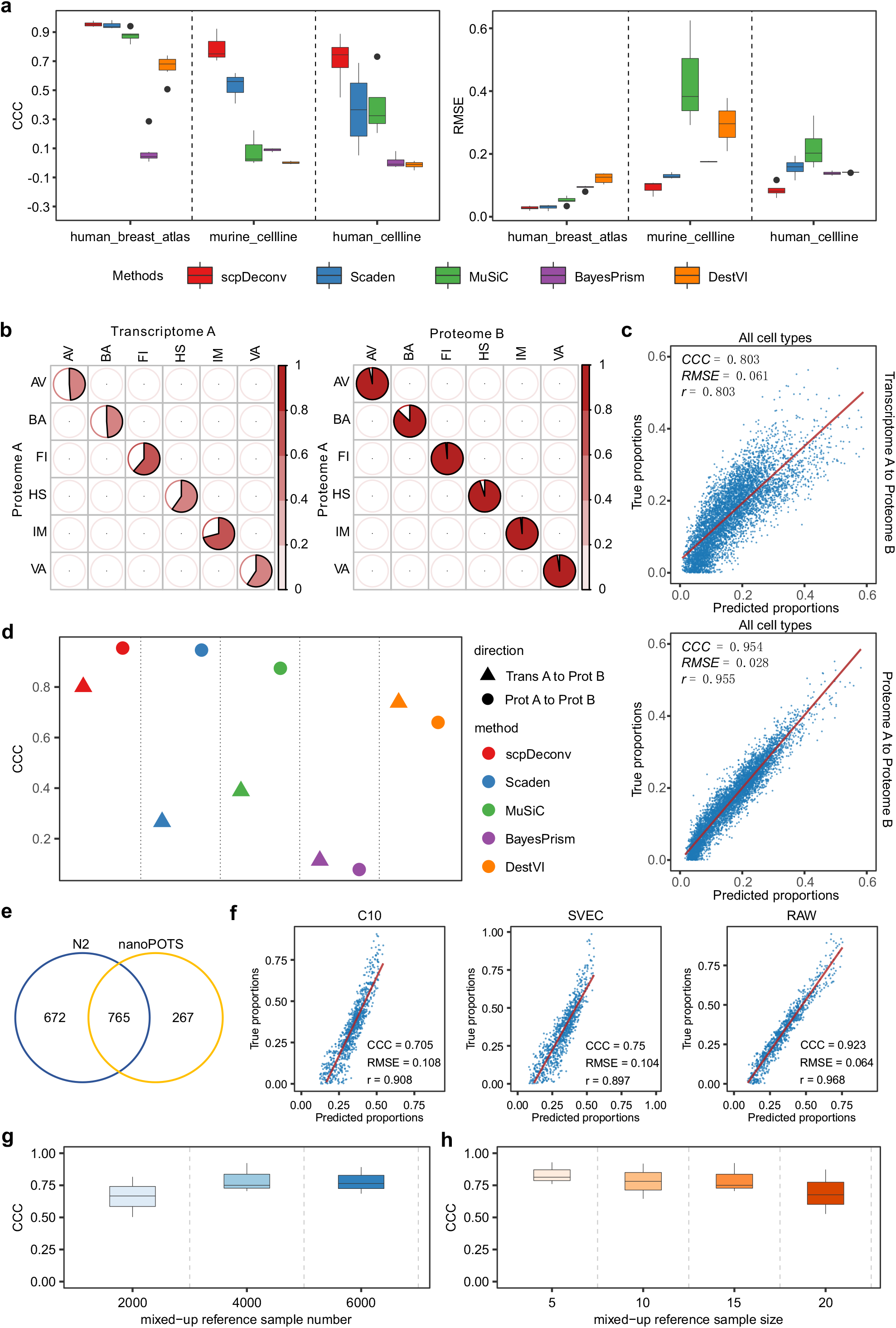
Overall performance comparation of cell type mixture deconvolution. a, CCC (left) and RMSE (right) values of scpDeconv, Scaden, MuSiC, BayesPrism, and DestVI on the testing cases of the human breast atlas datasets (proteome to proteome), murine cell line datasets, and human cell line datasets. Boxplots show the median (center lines), interquartile range (hinges), and 1.5 times the interquartile range (whiskers). b, Correlation plots between donor A’s proteome and donor A’s transcriptome (left) or donor A’s proteome and donor B’s proteome (right) within 6 cell types studied in human breast atlas datasets. AV: alveolar cells, HS: hormone sensing cells, BA: basal cells, FI: fibroblasts, VA: vascular lymphatic cells, IM: immune cells. c, Scatter plots of true (y axis) and predicted cell type proportions (x axis) by deconvolving pseudo-bulk proteomic data from donor B with single-cell transcriptomic data from donor A as reference (top) or with single-cell proteomic data from donor A as reference (bottom) in the case of human breast atlas datasets. d, CCC values of scpDeconv and the comparison methods by deconvolving pseudo-bulk proteomic data from donor B with single-cell transcriptomic data from donor A as reference (triangle) or with single-cell proteomic data from donor A as reference (circle) in the case of human breast atlas datasets. e, Venn diagram showing the overlap of proteins between two murine cell line datasets (N2 dataset and nanoPOTS dataset). f, Scatter plots of true (y axis) and predicted proportions (x axis) of three cell types (C10, SVEC, and RAW) in the case of murine cell line datasets using scpDeconv. g, CCC values of scpDeconv in the case of murine cell line datasets with setting different reference sample numbers in stage 1. h, CCC values of scpDeconv in the case of murine cell line datasets with setting different reference sample sizes in stage 1.

To compare the performance of prior cell type deconvolution algorithms with scpDeconv with transcriptomic and proteomic data, respectively, we carried out a deep exploration on the human breast atlas data. This is a large-scale clinical dataset where transcript data was generated by scRNA-seq and proteomic data was generated by CyTOF from 6 kinds of cell types present in breast tissue. Specifically, the cell types were alveolar cells, hormone sensing cells, basal cells, fibroblasts, vascular lymphatic cells, and immune cells^31^. 33 genes/proteins were measured in both the transcriptomic and proteomic data. To carry out the comparison under strict conditions, paired scRNA-seq data^(A)^ and CyTOF data^(A)^ from the same donor (A) and CyTOF data^(B)^ from another donor(B) were selected from the human breast atlas dataset. We extracted the mean expression/abundance of 33 genes/proteins and calculated the correlations between gene expression and protein abundances within each cell type. As shown in Figure 2b and Extended Data Figure 1b, the corresponding transcript (scRNA-seq data^(A)^) and protein (CyTOF data^(A)^) patterns showed relatively low correlation, even though they were sampled from the same donor. In contrast, the proteomic profiles from CyTOF data^(A)^ or CyTOF data^(B)^ were highly similar. Furthermore, to explore the benefits of using single cell proteomic data as reference in deconvolving bulk proteomic data, we simulated pseudo-bulk samples by using CyTOF data^(B)^, and then used scpDeconv to deconvolve them under the guidance of scRNA-seq data^(A)^ or CyTOF data^(A)^. In this case, compared with other methods, scpDeconv obtained the best performance on both deconvolution tasks with different reference data (Figure 2c-d and Extended Data Figure 1c). In addition, scpDeconv, Scaden and MuSiC performed better on the deconvolution task with CyTOF data^(A)^ as reference than that with scRNA-seq data^(A)^ as reference. In contrast, DestVI performed better on transcript data compared to CyTOF data due to its heavy reliance on the assumed distribution of transcripts.

Overall, these results indicate the necessity and the benefits of deconvolving bulk proteomic data with single-cell proteomic data as reference (Figure 2d and Extended Data Figure 1c).

In the case of the murine cell line datasets, we collected two independent single cell proteomic datasets using two advanced technologies (N2 and nanoPOTs) from the 3 common murine cell types (C10, RAW, and SVEC, respectively) grown under standard culture conditions. The two datasets showed a small overlap in protein types (Figure 2e) and different data distributions in terms of protein abundance (Extended Data Figure 1d). To assess scpDeconv’s performance on samples from different acquisition methods, we specified the mixed-up N2 dataset as reference data and the mixed-up nanoPOTS dataset as target data. We then executed scpDeconv to estimate proportions of C10, RAW, and SVEC for target data (Figure 2f) and compared the results with those generated by other deconvolution methods. In this test, scpDeconv achieved the best CCC score, RMSE and Pearson’s correlation values (Figure 2a, Extended Data Figure 1a). Furthermore, when altering the sample numbers/sizes of the mixed-up N2 dataset in stage 1, scpDeconv maintained stable performance and showed a high level of robustness (Figure 2g-h). In addition, we also validated the effect of stage 2 (imputation module) by removing it from the workflow of scpDeconv. This significantly lowered the performance on the deconvolution task (Extended Data Figure 1g).

### Performance of imputing missing proteins in cell type deconvolution on human cell line datasets

As mentioned in *Method* part, *Stage 2* of scpDeconv is specifically designed for proteomics deconvolution by developing an autoencoder that imputes missing proteins of reference data (especially the highly variable proteins observed in target data). To systematically analyze the effect of the imputation module on spcDeconv, we carried out an ablation study on missing proteins using the human cell line datasets by changing the extent of missing proteins in reference data. To construct a more complex scenario with large gaps of detected proteins and altered data distributions, we integrated 3 single cell human cell lines datasets (namely SCoPE2_Leduc, pSCoPE_Huffman, and pSCoPE_Leduc) as the reference group (Figure 3a), and 2 additional single cell human cell lines datasets (namely T-SCP and plexDIA) as the target group (Figure 3b). The datasets shared 4 human cell lines but suffered from severe batch effects and strikingly different identified protein types (Figure 3c, Extended Data Figure 2a). In particular, in the reference group, the highly variable proteins (HVPs) specific in the target group were not detected, although they are some extent informative for deconvolution task (Figure 3d and Extended Data Figure 2b). The scpDeconv imputation module in *Stage 2* was added to tackle this problem.

**Figure 3.**
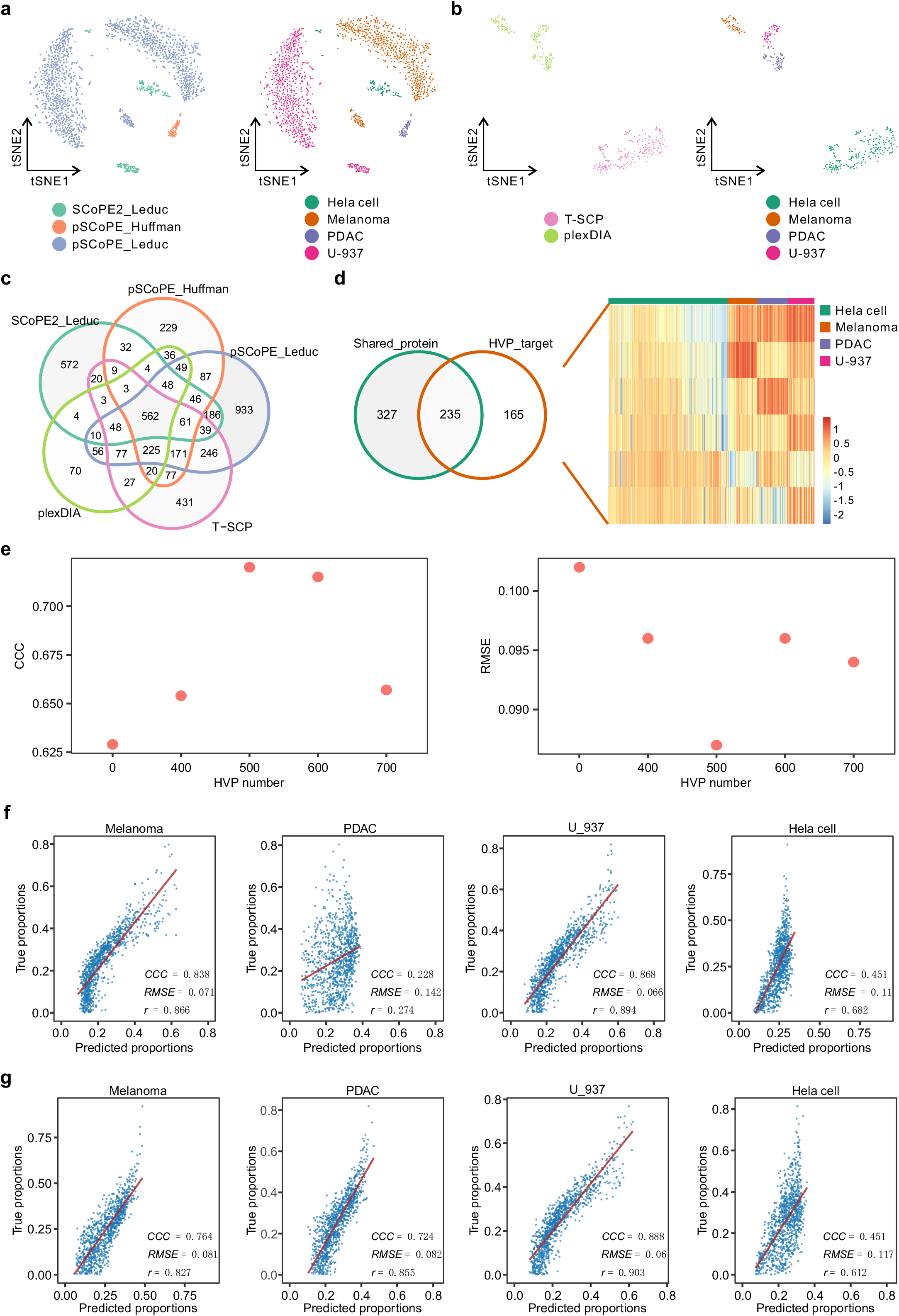
Deconvolution performance on integrated human cell line datasets with different HVP numbers. a, Integrated t-SNE plots colored by data sources (left panel) or cell types (right panel) in the reference group (SCoPE2_Leduc, pSCoPE_Huffman, and pSCoPE_Leduc datasets). b, Integrated t-SNE plots colored by data sources (left panel) or cell types (right panel) in the target group (T-SCP and plexDIA datasets). c, Venn diagram showing the overlap of proteins from five different human cell line datasets. d, Venn diagram showing the overlap between shared proteins in all datasets and highly variable proteins (HVPs) identified in target group. Heatmap showing the protein abundance profiles of specific HVPs in the target group. e, CCC (left panel) and RMSE values (right panel) of scpDeconv with different HVP numbers identified in the target group of human cell line datasets. f, Scatter plots of true (y axis) and predicted proportions (x axis) of four cell lines (Melanoma, PDAC, U-937, and Hela cell) in human cell line datasets using scpDeconv without considering target specific HVPs. g, Scatter plots of true (y axis) and predicted proportions (x axis) of four cell lines (Melanoma, PDAC, U-937, and Hela cell) in human cell line datasets using scpDeconv with setting target HVP number as 500.

The ablation study on missing proteins showed that the imputation module improved the model’s performance to some extent by supplementing input features with the undetected, target-specific HVPs. Performance improvement was particularly noticeable for the prediction accuracy of the proportion of PDAC cells in the sample (Figure 3e-g). In addition, we also explored the magnitude of the effect of target HVPs on the deconvolution task by varying the number of target HVPs between 400 to 700. The results showed that irrespective of the number of HVPs added the system achieved a better performance compared to the performance without target-specific HVPs (Figure 3e). As shown in Extended data figure 2c-e, after the imputation stage, missing features within the reference data can be complemented on the basis of target data.

### Quantitative evaluation of cell cycle deconvolution across cell lines

Besides the cell type deconvolution task, inference of cellular state is also critical for studying biological processes, tumor microenvironment, and cancer therapy^32–34^. Thanks to the ultra-high sensitivity of mass spectrometry, proteomic technologies can detect the subtle changes of functional proteins that determine or correlate with the state of a cell. This is exemplified by different cell cycle states which are characterized by well understood changes in abundance of multiple proteins^35,36^. To quantitatively evaluate scpDeconv’s performance for the challenging estimation of the distribution of cell cycle states in bulk proteomic samples, we collected bulk proteomic data with constant and pure cell cycle state and cell type for each sample (namely G1/S/G2-state melanoma, or G1/S/G2-state monocyte samples). We used these data to simulate pseudo-bulk proteomic data with randomly mixed-up cell states at a constant cell type distribution. Furthermore, to force the model to focus on cell cycle states rather than cell types, only functional proteins (top 20 for each state) corresponding to certain cell cycle states were chosen as input features (Figure 4a-b). By comparing the abundance of these 60 functional proteins between G1/S/G2-state melanoma and G1/S/G2-state monocyte samples, we noticed that functional proteins of cell cycle states were highly conserved between melanoma and monocyte samples (Figure 4c), which means that cell cycle deconvolution across cell types/lines is feasible. Therefore, we used scpDeconv, along with other deconvolution methods to compare their respective performance in estimating cell cycle state proportions for pseudo-bulk data simulated by G1/S/G2-state monocyte samples with G1/S/G2-state melanoma samples as reference (marked as *mel_to_mon*), and vice versa (marked as *mon_to_mel*). As shown in Figure 4d-e, and Extended Data Figure 3-4, compared with other deconvolution methods, scpDeconv achieved superior performance for both cell cycle state deconvolution studies, demonstrating the great potential of the new method for the inference of subtle differences in the distribution of cell cycle states in bulk samples and, by implication, the inference of other cell cycle states, provided proteins characteristic for these states are known and detectable.

**Figure 4.**
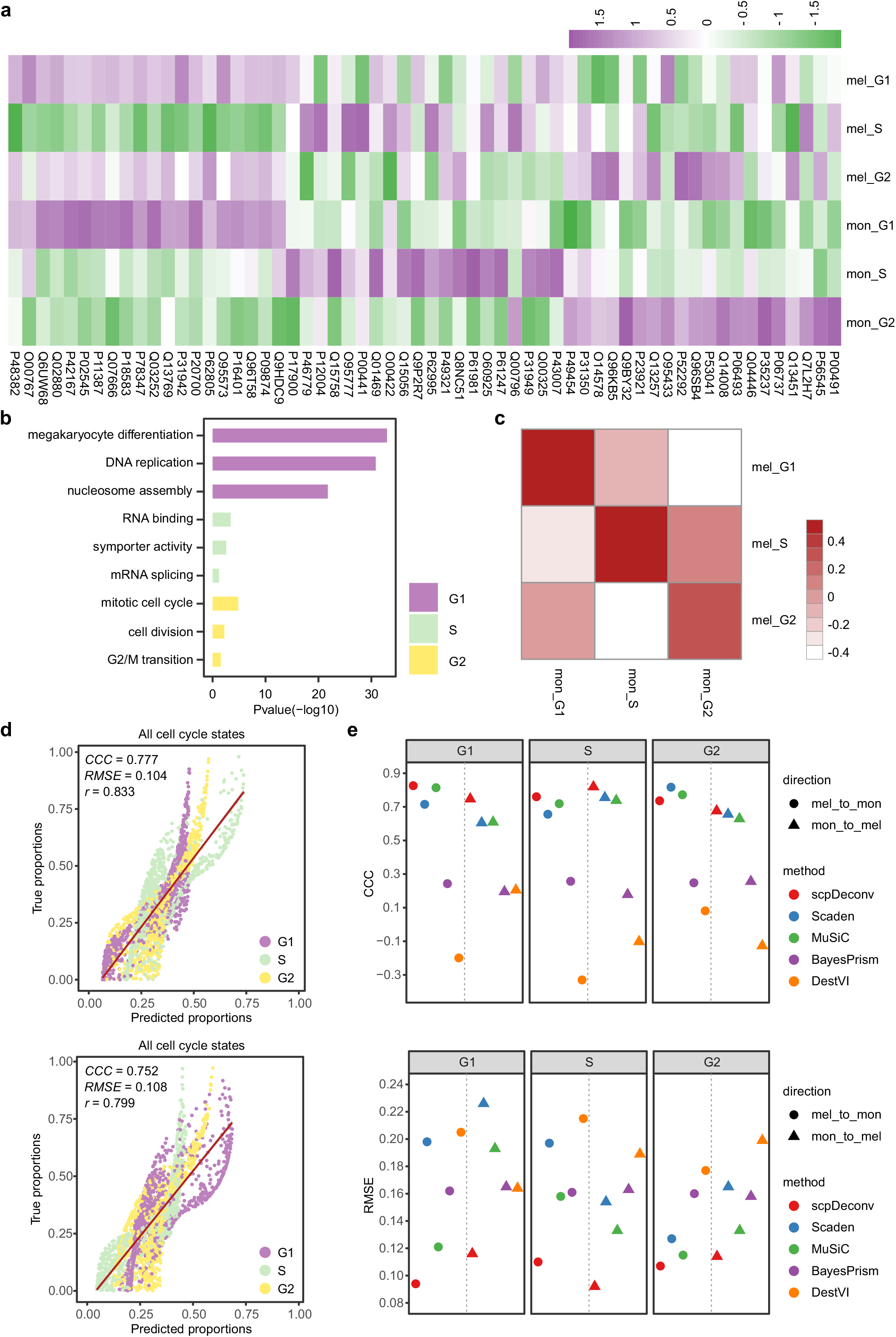
Inference of cell cycle across cell lines. a, Heatmap of the abundance of cell cycle specific marker proteins in melanoma or monocyte samples. b, GO enrichment of marker proteins of G1, S, or G2 phases. c, Heatmap showing the correlation of marker proteins’ abundance in melanoma and monocyte samples. d, Scatter plots of true (y axis) and predicted proportions (x axis) of three cell cycle states (G1, S, and G2) by deconvolving monocytes (mon) pseudo samples with melanoma (mel) sample as reference (top) and vise versa (bottom) using scpDeconv. e, CCC (top) and RMSE (bottom) values of scpDeconv and the comparison methods by deconvolving monocytes pseudo samples with melanoma samples as reference (circle) and vise versa (triangle).

### Application to clinical tissue proteomics

Melanoma is a rare but invasive cancer of the skin, with high rates of growth and metastasis^37^. Previous clinical proteomic analyses of melanoma samples have focused on the detection of proteins as markers for disease stratification and responsiveness to therapy. Generally, these studies have not considered the tumor microenvironment and the effect of cellular heterogeneity on clinical outcomes^38^. Here, we explored the capacity of scpDeconv to re-analyze representative clinical proteomic data from the bulk analysis of 174 metastatic melanoma Grade IV samples consisting of 4884 high quality protein identifications^39^. As shown in Figure 5a and Extended Data Figure 5a, scpDeconv used the most advanced spatial proteomics (Deep Visual Proteomics, DVP) on melanoma tissues as reference data, which contained 7 cell classes (vertical growth melanoma, radial growth melanoma, CD146-high melanoma, CD146-low melanoma, in situ melanoma, melanocytes, and stroma) identified in the tumor microenvironment^40^. Following the procedure of scpDeconv, the proportions of 7 cell classes were estimated from the tissue proteomic profiles of 174 melanoma samples (Figure 5b-c). The results indicated that the vertical growth melanoma cell class was dominant in the late-stage metastasis melanoma samples, while in situ melanoma cell class was the minority, consistent with previous clinical research and indicating that the presence of vertical growth phase signifies that the melanoma acquires the capacity of metastasis^41^. We further performed survival analysis using the predicted cell class proportions for 174 melanoma samples as input data (Figure 5d-f, Extended Data Figure 5b-e). Compared with other cell classes, the proportion of vertical growth melanoma and radial growth melanoma exhibited significant indications for survival risk of melanoma patients. The patients with higher vertical growth melanoma proportions tended to gain lower survival probability, while those with higher radial growth melanoma proportions tended to gain higher survival probability. Consistent with previous studies, these results showed that in contrast to melanoma in radial growth stage, melanoma with vertical growth pattern usually accompanies with strong aggressivity and poor prognosis^42,43^.

**Figure 5.**
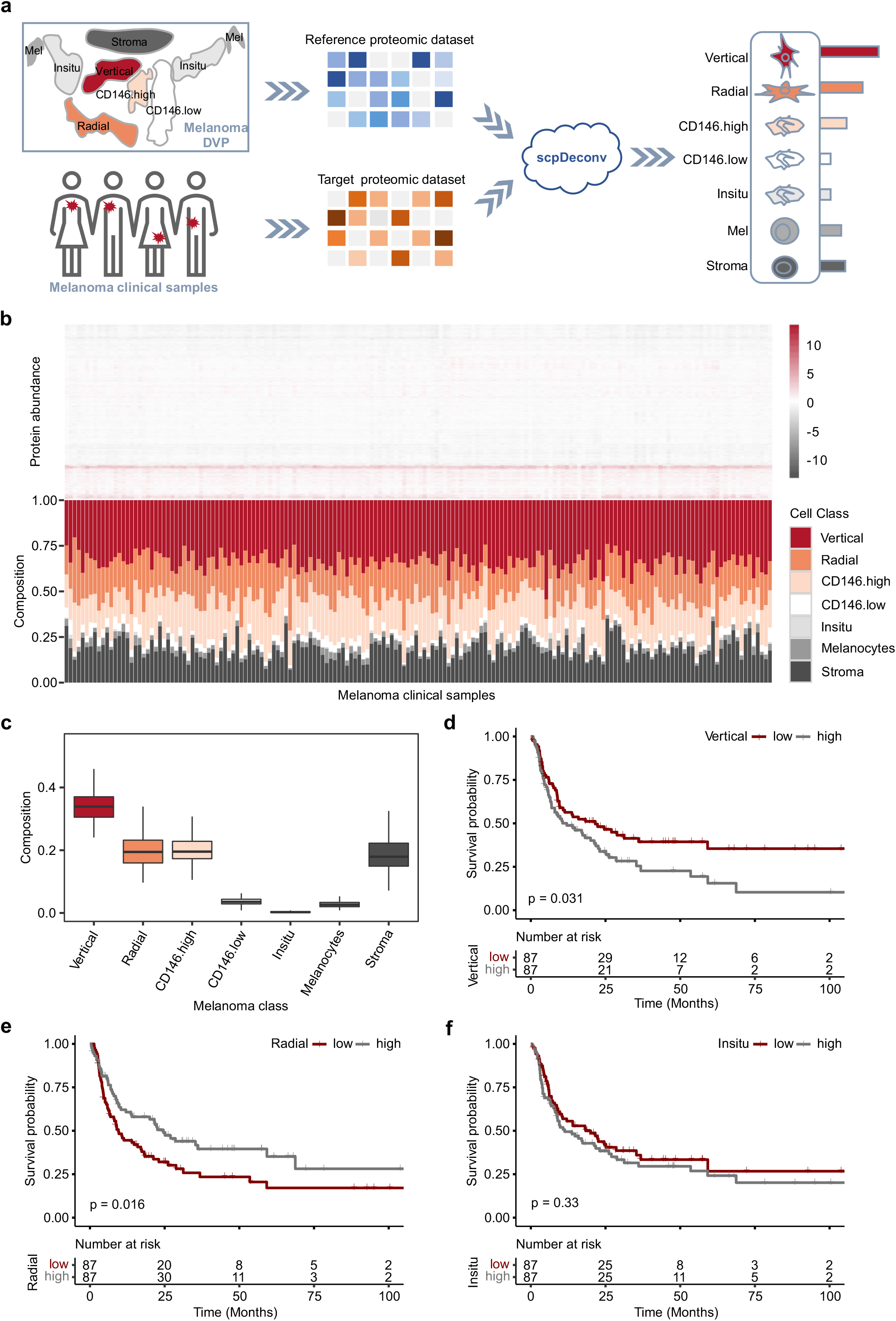
Application on clinical tissue proteomic data of metastasis melanoma. a, The pipeline of applying scpDeconv on deconvolving clinical tissue proteome of metastasis melanoma patients with melanoma DVP dataset as reference. b, Heatmap of proteomic profile (top) and histogram of predicted cell class proportions (bottom) for each sample in clinical melanoma dataset. c, Distribution of predicted cell class proportions of clinical melanoma samples. Box plots show the median (center lines), interquartile range (hinges), and 1.5 times the interquartile range (whiskers). d, KM plot of the predicted proportions of vertical growth melanoma for 174 melanoma patients. e, KM plot of the predicted proportions of radial growth melanoma for 174 melanoma patients. f, KM plot of the predicted proportions of in situ melanoma for 174 melanoma patients.

In addition, as mutation of the BRAF protein is reported to reflect the tumorigenesis of melanoma, we further combined the indication of predicted cell class proportions with the occurrence of BRAF mutation (Extended Data Figure 6a). In this cohort, we can observe that BRAF mutation’s indicator power is much weaker than our predicted vertical/radial growth melanoma proportions (Extended Data Figure 6b). By grouping samples into BRAF^(+)^ (mutation) and BRAF^(-)^ (non-mutation), the predicted vertical or radial growth melanoma proportion showed more significant prognostic indication in BRAF^(+)^ samples (Extended Data Figure 6c-f).

Overall, the application of scpDeconv on a clinical proteome cohort indicated that the tumor microenvironment could be reflected by predicted cell type proportions and demonstrated its potential to be a valuable tool for cancer prognosis.

## Discussion

Single-cell proteomic technology is rapidly developing and is becoming an important component of biological and clinical research^25^. One particularly important use case is the deconvolution of cell types/states from bulk tissue proteome profiling data, a task that has not yet been achieved. By fully considering the inherent characteristics of the recently developed single-cell proteomics technologies, we developed a novel deconvolution method, scpDeconv, that is specifically tailored to proteomic data. By feat of an autoencoder network, scpDeconv’s imputation module can impute values for low-abundance proteins missing in single-cell proteomic data. This is achieved by joint-learning on both, mixed-up single-cell proteomic data and target bulk proteomic data. Through domain-adversary training, scpDeconv can extract the domain-invariant latent features of imputed reference data and transfer labels to target data. Rigorous and extensive experiments evaluated scpDeconv’s superior performance and robustness on cell type deconvolution as well as the inference of cell cycle states from bulk tissue proteomic datasets. Furthermore, re-analysis of an existing bulk proteomic dataset from clinical melanoma samples showed the clinical application value of scpDeconv in cancer diagnosis and prognosis. Compared to the existing deconvolution methods tailored to transcriptomics, scpDeconv offers four levels of advantages for proteome data deconvolution: i), scpDeconv infers cell type proportions based on joint-learning of reference data and target data in a data-driven manner and is thereby unrestricted by the prior assumptions of data distribution and compatible with a broad range of proteomic data from different experiments or technology background; ii), scpDeconv introduces a domain adaptation strategy which can efficiently handle batch effects and close the distribution gaps between reference data and target data; iii), scpDeconv can impute missing values of low-abundance proteins in single cell proteomic data and improve the quality of reference data with the help of an imputation module; iv), we have validated that the output of scpDeconv is capable of reflecting cell type distribution in the tumor microenvironment and that the thus identified cell type distributions are useful for disease diagnosis and prognosis. At present, scpDeconv shows some limitations. Reference proteomic data with known cell type composition needed for model training are constructed from single-cell proteomic data with labelled cell types. At present single-cell proteomics is still in its infancy and has not been applied to a broad range of biological samples. The single-cell proteomic data we exhaustively collected in this study only involve a minority of cell types, thereby limiting our method’s applicability at present.

This first algorithm tailored for cell type/state deconvolution from proteomic data using single-cell proteomic data as reference will have numerous key applications. These include the deconvolution of cell type proportions, estimation of cell states, exemplified by cell cycle states, and re-analysis of the microenvironment of multiple cancers thus adding additional value to already acquired bulk proteomic datasets. We therefore anticipate that scpDeconv will benefit the proteomics community by supporting the extraction of additional, biologically and clinically important results from bulk proteomic datasets. With the exploration of the publicly available single-cell proteomic atlas, it will be feasible in the future to re-analyse existing tissue proteomic data within the broader context. Furthermore, with a set of reliable microenvironment references of a vast range of cancer types, we believe that scpDeconv will be capable of analysing all the components of certain tumor samples at low cost and benefiting for basic medical research.

## Methods

### The model

The overall model architecture is shown in Figure 1, containing 4 stages: i), reference data preparation; ii), missing features imputation and data quality improvement; iii), domain-adversarial training and deconvolution; iv), cell type proportion inference for target samples.

#### Stage 1. Reference data preparation

As a supervised deconvolution method, single-cell proteomic data labelled with cell types are needed to prepare reference data with known cell type fractions. And the bulk proteomic dataset from similar cohort is termed the target data to be deconvolved. The reference data and target data are usually sourced from different sequencing platforms and experiments, thereby having gaps in data distribution and proteome coverage. Bulk proteomics can usually detect much more proteins compared to single-cell proteomics, in which the low-abundance proteins are hard to detect.

##### Mix-up strategy

Mix-up is an efficient way of data augmentation by a random combination of inputs^44^. Here we employed it on single-cell proteomic data to make up pseudo-bulk proteomic data before training. The random combination of cell type distributions from single-cell proteomic data increases the data diversity of the reference dataset for training, thereby improving the model’s generalizability. In this stage, we generated the pseudo bulk proteomic data 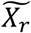 with random cell type proportions 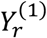 as reference data, and then sent it to the autoencoder module of stage 2 for data imputation.

##### Data alignment

Before model training, we need to align proteins of reference data and target data for model input. We firstly extracted the proteins shared between reference and target data. And then we took the union set of these shared proteins and HVPs specific in target data as input feature set, since the target data are supposed to contain more proteins than the reference data, and those target-specific HVPs may be informative for deconvolution task. The missing proteins in the input feature set for reference data are initialized to zeros and will be imputed in *Stage 2*.

#### Stage 2. Missing features imputation and data quality improvement

Due to the immaturity of single-cell proteomic technology, the reference data generated by single cell proteomic data have strong noises, large-scale missing low-abundance proteins, and an obvious distribution gap with the target bulk proteomic data. To compensate for the above obstacles caused by single-cell proteomic technology, we employed an autoencoder module to close the gap between the reference data and the target data, impute missing proteins of the reference data, and improve the quality of reference data. As shown in Figure 1b, the target data and the reference data generated in *Stage 1* are inputted into Encoder1. And then the latent embedding generated by Encoder1 is sent to a decoder to reconstruct the input. The loss between the original input and the reconstructed output is MSE loss. The latent embedding is also sent to a deconvolution head to estimate cell type proportions of reference data, and the L1 loss is calculated between the predictions and its known proportions. The formulations of the architecture in *Stage 2* are:

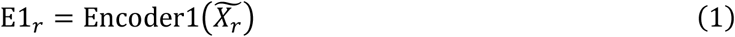

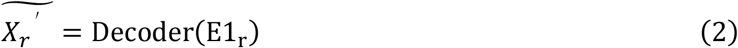

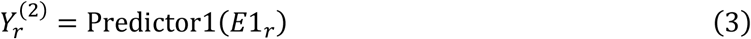

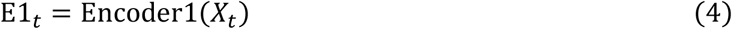

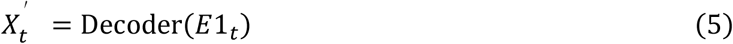

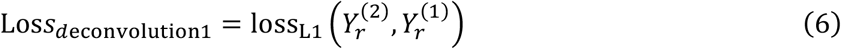

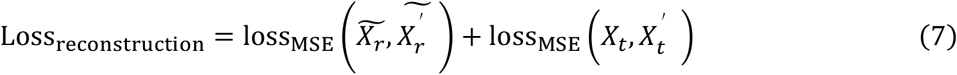

Where 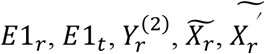, *X*_*t*_ and 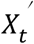 are the embedding of the reference data, the embedding of the target data, the output of the Predictor1 used for deconvolution, the reference proteomic data, the output of decoder with the embedding of reference data, the target proteomic data, and the output of decoder with the embedding of the target data, respectively. After the convergence of the training stage, the imputed reference data 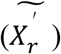 are used as the input of *Stage 3*.

#### Stage 3. Domain-adversarial training and deconvolution

As shown in Figure 1c, the imputed reference data from *Stage 2* and the target data from *Stage 1* are sent to the Encoder2 in *Stage 3*, and the encoded features are sent to the domain discriminator and the deconvolution head. The domain discriminator is employed to discriminate whether the latent features are encoded from the reference data or the target data. By jointly training the Encoder2 and the discriminator in this *stage*, the Encoder2 is prone to extracting domain-invariant features, which is suitable for the deconvolution task in which the reference data and the target data are from different sources. The deconvolution head consists of 2 fully connected layers and is used for predicting cell type proportions of target data. The Encoder2 and deconvolution head are jointly trained with the imputed reference data as input and the known proportions as supervision signals. The two kinds of joint-training (Encoder2 and discriminator/Encoder2 and deconvolution head) are then executed iteratively until both losses are converged.

As for domain adaptation, we calculated BCE loss and optimized our model’s weights by minimizing the distributions between the reference and target domains. The formulations of the architecture in *Stage 3* are:

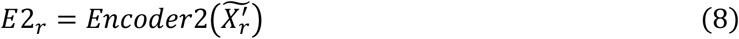

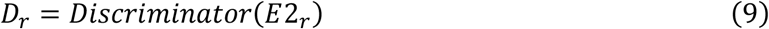

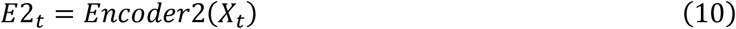

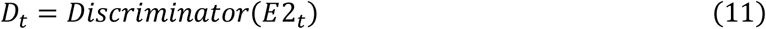

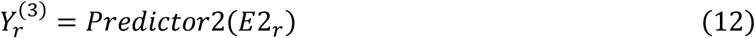

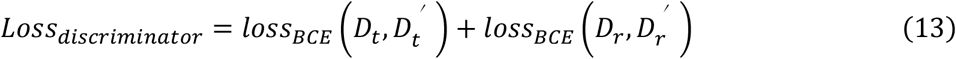

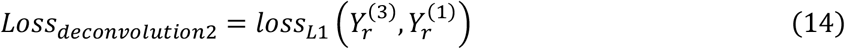

Where 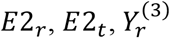, *D*_*r*_ and *D*_*t*_ are the embedding of imputed reference data by Encoder2, the embedding of the target data by Encoder2, the output of the deconvolution head, the output of discriminator with the embedding of imputed reference data, and the output of discriminator with the embedding of target data, respectively.

#### Stage 4. Cell type proportion inference for target samples

As shown in Figure 1d, after domain-adversary training in *Stage 3*, the trained Encoder2 and deconvolution head are used to infer the cell type proportions of target samples. The formulation of the architecture employed in model inference is:

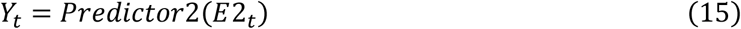

Where *E2*_*t*_ is the output in *Formula* (10) with *X*_*t*_ as input, and *Y*_*t*_ is the final predicted cell type proportions of the target data.

### Data

The datasets used in this study are listed in Supplemental Table 1.

#### Human breast atlas datasets

In the human breast atlas dataset^31^, scRNA-seq was performed on 16 breast tissues (52,681 cells) and CyTOF was performed on 38 breast tissues (751,970 cells,40 protein markers involved in breast development and tumorigenesis). The scRNA-seq and CyTOF datasets have 6 cell types in common (alveolar cells, hormone sensing cells, basal cells, fibroblasts, vascular lymphatic cells, and immune cells). The processed scRNA-seq dataset was downloaded from GEO (accession no. GSE180878) and the processed CyTOF dataset was downloaded from https://data.mendeley.com/datasets/vs8m5gkyfn/1.

#### Murine cell line datasets

The murine cell line datasets come from two single-cell proteomic technologies, N2 and nanoPOTS, applied in 3 identical murine cell lines (RAW 264.7, a macrophage cell line; C10, a respiratory epithelial cell line; SVEC, an endothelial cell line)^24,49^. N2 dataset (Murine cell line dataset generated by N2) includes 108 cells (36 RAW cells,36 C10 cells, and 36 SVEC cells) with 1,437 high quality protein identifications, and was downloaded from MassIVE data repository (accession no. MSV000086809). nanoPOTS dataset (Murine cell line dataset generated by nanoPOTS) includes 72 cells (24 RAW cells,24 C10 cells, and 24 SVEC cells) with 1,032 high quality protein identifications, and was downloaded from MassIVE data repository (accession no. MSV000084110). These two datasets have 762 proteins in common.

#### Human cell line datasets

The human cell line datasets consist of five different single cell proteomic datasets, namely T-SCP, plexDIA, pSCoPE_Leduc, SCoPE2_Leduc, and pSCoPE_Huffman, generated by different state-of-the-art technologies^36,45–48^. T-SCP dataset contains 434 Hela cells and 2501 proteins and was downloaded from PRIDE partner repository (accession no. PXD024043). plexDIA was performed on 155 Melanoma, PDAC, and U-937 cells and detected 1475 high quality protein identifications, and was downloaded from https://scp.slavovlab.net/Derks_et_al_2022. pSCoPE_Leduc dataset (involving 1755 Melanoma and U-937 cells and 2844 proteins) was generated by pSCoPE technology and was downloaded from https://scp.slavovlab.net/Leduc_et_al_2022. SCoPE2_Leduc dataset (involving 163 Hela cell and U-937 cell and 1647 proteins) was generated by SCoPE2 technology and was downloaded from https://scp.slavovlab.net/Leduc_et_al_2021. pSCoPE_Huffman dataset (involving 206 PDAC cells and 1659 proteins) was also generated by pSCoPE technology and was downloaded from https://scp.slavovlab.net/Huffman_et_al_2022. After integration, these five datasets involved 4 cell types in total, consisting of 1,755 melanoma,434 Hela cells,579 PDAC cells, and 163 U-937.

#### Bulk proteomic cell cycle datasets

Bulk proteomic datasets in which each sample has pure and constant cell type and state (G1/S/G2-state melanoma and G1/S/G2-state monocyte samples) were generated in the same experiment as pSCoPE_Leduc dataset^48^. And following the authors’ reported procedure, the top 20 highly expressed marker proteins of each cell cycle state (G1, S, G2) were selected as model’s input features.

#### Deep Visual Proteomics dataset

Deep Visual Proteomics (DVP) dataset links protein abundance of melanoma samples to complex cellular or subcellular phenotypes while preserving spatial context, involving average 1910 proteins^40^. This dataset contains 7 cell classes (vertical growth melanoma, radial growth melanoma, CD146-high melanoma, CD146-low melanoma, in situ melanoma, melanocytes, and stroma) within the tumor microenvironment of melanoma, defined by spatial location and SOX10/CD146 staining intensity. We downloaded this dataset from the PRIDE partner repository (accession no. PXD023904).

#### The clinical proteomic dataset of metastatic melanoma

The bulk proteomic dataset of metastatic melanoma was collected from the previous clinical study, in which mass spectrometry-based proteomic analysis was performed in 174 metastasis melanoma patients, with 4883 high quality protein identifications^50^.

### Training details

For all of the cases, the batch size of training process in stage 2 and stage 3 was set to 50, and the learning rate was set to 0.0001. The other training details, such as pseudo-sample number and pseudo-sample size (cell number in each pseudo-sample) of the reference data or target data, the number of HVPs identified in the target data, and the number of training epochs used in each case can be found in Supplemental Table 2.

Furthermore, in the case of murine cell line datasets, to explore scpDeconv’s stability, we set different reference sample numbers (2000,4000, and 6000) with fixed sample size, and different reference sample sizes (5,10,15, and 20) with fixed sample number.

### Comparison of methods

#### Scaden

Scaden is a deep-learning-based deconvolution algorithm that uses RNA-seq data and paired scRNA-seq data to infer the cellular composition of tissues^9^. The basic architecture of Scaden is a neural network with 4 fully connected layers which takes the pseudo bulk RNA-seq data as input and the predicted cell type fractions as output, using L1 loss during training to minimize the error between the predicted and real cell type fractions. The final Scaden model is ensembled by three best-performing models with different parameter combinations.

#### MuSiC

MuSiC uses cell-type specific gene expression from scRNA-seq data to deconvolve bulk RNA-seq data in tissues^4^. MuSiC first reweighs genes based on cross-subject variance (informativeness), and then uses a recursively clustering strategy to gather related cell types. The proportions of cell types are estimated by non-negative least squares regression based on a series of assumptions.

#### BayesPrism

BayesPrism is a Bayesian-based deconvolution method for cancer tissues depending on patient-derived scRNA-seq as prior information^29^. The Bayesian algorithm models prior distribution learned from the paired scRNA-seq data and then infers joint posterior distribution of cell type proportions as the deconvolution results.

#### DestVI

DestVI is also a Bayesian-based method that learns cell type specific profiles and latent variations using a conditional deep generative model^30^. The method assumes that transcripts profiles follow a negative binomial distribution to model the reference scRNA-seq data and spatial transcriptomic data.

### Evaluation experiments

#### Cell type mixture decomposition with proteome profiling

As for testing the deconvolution performance with the reference data and target data from different modalities, in the simulation case of human breast atlas datasets, we deconvolved the simulated bulk proteomic data constructed from CyTOF data with scRNA-seq data in the same cohort as reference or with CyTOF data in the same cohort as reference, splitting the dataset into reference and target data by donors.

There are apparent batch effects between different proteomic technologies and large gaps between the distributions of bulk proteomic data and single-cell proteomic data. Therefore, to evaluate scpDeconv’s deconvolution performance on the situation where the bulk/single-cell proteomic data came from totally different proteomic technologies and different experiments, we deconvolved bulk proteomic data simulated by nanoPOTS dataset with the mixed-up N2 dataset as reference.

Furthermore, we employed the integrated human cell line datasets to evaluate scpDeconv’s performance on integrated multi-centre datasets. The Melanoma, PDAC, U-937, and Hela cell lines appeared in different experiments were selected for testing.

#### Systematic analysis

To explore the efficiency of the imputation of missing proteins in *Stage 2*, we executed cell type deconvolution experiments on the integrated human cell line datasets with considering target-specific HVPs or without considering target-specific HVPs, and also compared deconvolution performance between model with imputation stage and model without imputation stage. For the experiments without considering target-specific HVPs, we only used the overlap proteins from both reference and target datasets as input features. For the experiments with considering target-specific HVPs, we changed the number of HVPs identified in target data from 400 to 700.

### Application

#### Cell cycle inference with proteome profiling

The cell cycle states reflect the replication process of cancer cells and can be represented by the subtle changes of cell-cycle-relevant proteins. We collected the bulk proteomic data of melanoma or monocyte samples with pure cell subpopulations within G1, S, or G2 stage. Then the melanoma cell cycle mixtures were constructed by mixing up the melanoma cell subpopulations within G1, S, or G2 stage, and monocyte cell cycle mixtures were constructed by mixing up the monocyte subpopulations within G1, S, or G2 stage. Furthermore, to estimate scpDeconv’s performance on the cell cycle inference task across cell types/cell lines, we deconvolved the melanoma cell cycle mixtures with monocyte cell cycle profiles as reference, and also deconvolved the monocyte cell cycle mixtures with melanoma cell cycle profiles as reference.

#### Investigation of microenvironments and clinical outcomes with clinical tissue proteome profiling

The establishment of scpDeconv that is tailored for cell type deconvolution of proteomic data opens up an opportunity to mine the tumor microenvironments of previously released clinical tissues with provided bulk proteomic data. We used scpDeconv to deconvolve the clinical proteomic data from metastasis melanoma cohort and assessed the impact of tumor class proportions on the prognostic determination according the survival analysis.

### Evaluation metrics

The linear concordance between ground truth ratios and predicted proportions of cell types or cell states is strictly evaluated by 3 metrics, including Lin’s concordance correlation coefficient (CCC), Root Mean Square Error (RMSE), and Person product-moment correlation coefficient (*r*). The formulas are as follows:

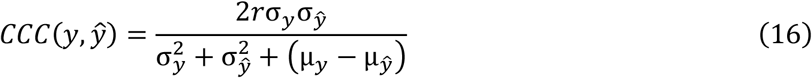

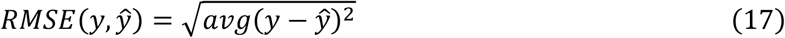

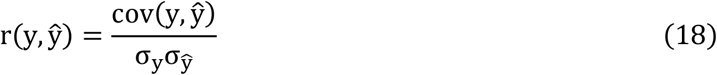

Where *y* are the ground truth cell type (or cell cycle state) proportions, *ŷ* are the prediction of cell type (or cell cycle state) proportions, *σ*_*y*_ is the standard deviation of *y, σ* _*ŷ*_ is the standard deviation of *ŷ, cov(y, ŷ)* is the covariance of *y* and *ŷ*, and µ_*y*_, µ_*ŷ*_ respectively are the mean of *y* and *ŷ*.

### Statistical analysis

Kaplan-Meier survival analysis is conducted to assess if the predicted cell type proportions are associated to outcomes (overall survival) in the clinical cohort. The P values are calculated using the log-rank test and are considered statistically significant if they are two-tailed less than 0.05.

### Biological analysis

Highly variable proteins are selected by Scanpy^51^. The Gene Ontology (GO) and pathway enrichment are performed using DAVID platform^52^.

## Data Availability

All data used in this study are publicly available and the usages were fully illustrated in the Method section.

## Code Availability

The codes will be released after the publication of this manuscript. The comparison methods are implemented under their official guidance and the reference codes are listed in the Method section.

## Acknowledgements

We thank Doctor Chen Li for giving suggestions for the project.

## Author Contributions

F.Y. and J.Y. conceived and designed the project. F.W. developed the algorithm and conducted experiments under the supervision of F.Y. and J.Y.. F.W. and F.Y. analysed the results. F.Y. and F.W. wrote the manuscript. F.W. finished the figures under the guidance of F.Y. and J.Y.. J.Y., G.W., R.A. and R.G. revised the manuscript. L.H. gave suggestions for the domain adaptation and the improvement of the manuscript. J.S. gave suggestions for the project. F.W. and F.Y. contributed equally. All authors reviewed and approved the manuscript.

## References

1. Galon, J. et al. Type, density, and location of immune cells within human colorectal tumors predict clinical outcome. Science (1979) 313, 1960–1964 (2006).

2. Ino, Y. et al. Immune cell infiltration as an indicator of the immune microenvironment of pancreatic cancer. British Journal of Cancer 2013 108:4 108, 914–923 (2013).

3. Shang, B., Liu, Y., Jiang, S. J. & Liu, Y. Prognostic value of tumor-infiltrating FoxP3+ regulatory T cells in cancers: a systematic review and meta-analysis. Scientific Reports 2015 5:1 5, 1–9 (2015).

4. Wang, X., Park, J., Susztak, K., Zhang, N. R. & Li, M. Bulk tissue cell type deconvolution with multi-subject single-cell expression reference. Nature Communications 2019 10:1 10, 1–9 (2019).

5. Qian, J. et al. A pan-cancer blueprint of the heterogeneous tumor microenvironment revealed by single-cell profiling. Cell Research 2020 30:9 30, 745–762 (2020).

6. Azizi, E. et al. Single-Cell Map of Diverse Immune Phenotypes in the Breast Tumor Microenvironment. Cell 174, 1293-1308.e36 (2018).

7. Frishberg, A. et al. Cell composition analysis of bulk genomics using single-cell data. Nature Methods 2019 16:4 16, 327–332 (2019).

8. Newman, A. M. et al. Determining cell type abundance and expression from bulk tissues with digital cytometry. Nature Biotechnology 2019 37:7 37, 773–782 (2019).

9. Menden, K. et al. Deep learning-based cell composition analysis from tissue expression profiles. Sci Adv 6, (2020).

10. Cable, D. M. et al. Robust decomposition of cell type mixtures in spatial transcriptomics. Nature Biotechnology 2021 40:4 40, 517–526 (2021).

11. Patrick, E. et al. Deconvolving the contributions of cell-type heterogeneity on cortical gene expression. PLoS Comput Biol 16, e1008120 (2020).

12. Lv, X. et al. Deconvolute Gene Expression Based on Deep Learning in Scrna-Seq. Proceedings - 2021 17th International Conference on Computational Intelligence and Security, CIS 2021 70–74 (2021) doi:10.1109/CIS54983.2021.00023.

13. Lin, Y. et al. DAISM-DNNXMBD: Highly accurate cell type proportion estimation with in silico data augmentation and deep neural networks. Patterns 3, 100440 (2022).

14. Edwards, N. J. et al. The CPTAC data portal: A resource for cancer proteomics research. J Proteome Res 14, 2707–2713 (2015).

15. Marguerat, S. et al. Quantitative Analysis of Fission Yeast Transcriptomes and Proteomes in Proliferating and Quiescent Cells. Cell 151, 671–683 (2012).

16. Gygi, S. P., Rochon, Y., Franza, B. R. & Aebersold, R. Correlation between Protein and mRNA Abundance in Yeast. Mol Cell Biol 19, 1720–1730 (1999).

17. Liu, Y., Beyer, A. & Aebersold, R. On the Dependency of Cellular Protein Levels on mRNA Abundance. Cell 165, 535–550 (2016).

18. Specht, H. & Slavov, N. Transformative Opportunities for Single-Cell Proteomics. J Proteome Res 17, 2565–2571 (2018).

19. Strack, R. Spatial proteomics with subcellular resolution. Nature Methods 2022 19:7 19, 780–780 (2022).

20. Petelski, A. A. et al. Multiplexed single-cell proteomics using SCoPE2. Nature Protocols 2021 16:12 16, 5398–5425 (2021).

21. Woo, J. et al. High-throughput and high-efficiency sample preparation for single-cell proteomics using a nested nanowell chip. Nature Communications 2021 12:1 12, 1–11 (2021).

22. Cheung, R. K. & Utz, P. J. CyTOF—the next generation of cell detection. Nature Reviews Rheumatology 2011 7:9 7, 502–503 (2011).

23. Slavov, N. Unpicking the proteome in single cells:: Single-cell mass spectrometry will help reveal mechanisms that underpin health and disease. Science 367, 512 (2020).

24. Woo, J. et al. High-throughput and high-efficiency sample preparation for single-cell proteomics using a nested nanowell chip. Nature Communications 2021 12:1 12, 1–11 (2021).

25. Perkel, J. M. Single-cell proteomics takes centre stage. Nature 597, 580–582 (2021).

26. Doerr, A. Single-cell proteomics. Nature Methods 2018 16:1 16, 20–20 (2018).

27. Vanderaa, C. & Gatto, L. Replication of single-cell proteomics data reveals important computational challenges. https://doi.org/10.1080/14789450.2021.1988571 18, 835–843 (2021).

28. Menden, K. et al. Deep learning–based cell composition analysis from tissue expression profiles. Sci Adv 6, (2020).

29. Chu, T., Wang, Z., Pe’er, D. & Danko, C. G. Cell type and gene expression deconvolution with BayesPrism enables Bayesian integrative analysis across bulk and single-cell RNA sequencing in oncology. Nature Cancer 2022 3:4 3, 505–517 (2022).

30. Lopez, R. et al. DestVI identifies continuums of cell types in spatial transcriptomics data. Nature Biotechnology 2022 40:9 40, 1360–1369 (2022).

31. Gray, G. K. et al. A human breast atlas integrating single-cell proteomics and transcriptomics. Dev Cell 57, 1400-1420.e7 (2022).

32. Warrener, R. et al. Tumor cell-specific cytotoxicity by targeting cell cycle checkpoints. The FASEB Journal 17, 1–21 (2003).

33. Li, J. & Stanger, B. Z. Cell Cycle Regulation Meets Tumor Immunosuppression. Trends Immunol 41, 859–863 (2020).

34. Schwartz, G. K. & Shah, M. A. Targeting the cell cycle: A new approach to cancer therapy. Journal of Clinical Oncology 23, 9408–9421 (2005).

35. Brunner, A.-D. et al. Ultra-high sensitivity mass spectrometry quantifies single-cell proteome changes upon perturbation. Mol Syst Biol 18, e10798 (2022).

36. Leduc, A., Huffman, R. G. & Slavov, N. Droplet sample preparation for single-cell proteomics applied to the cell cycle. doi:10.1101/2021.04.24.441211.

37. Balch, C. M. et al. Final Version of 2009 AJCC Melanoma Staging and Classification. Journal of Clinical Oncology 27, 6199 (2009).

38. Betancourt, L. H. et al. The Human Melanoma Proteome Atlas—Complementing the melanoma transcriptome. Clin Transl Med 11, e451 (2021).

39. Beck, L. et al. Clinical proteomics of metastatic melanoma reveals profiles of organ specificity and treatment resistance. Clinical Cancer Research 27, 2074–2086 (2021).

40. Mund, A. et al. Deep Visual Proteomics defines single-cell identity and heterogeneity. Nature Biotechnology 2022 40:8 40, 1231–1240 (2022).

41. Crowson, A. N., Magro, C. M. & Mihm, M. C. Prognosticators of melanoma, the melanoma report, and the sentinel lymph node. Modern Pathology 2006 19:2 19, S71–S87 (2006).

42. Prognosis and survival for melanoma skin cancer | Canadian Cancer Society. https://cancer.ca/en/cancer-information/cancer-types/skin-melanoma/prognosis-and-survival.

43. Ciarletta, P., Foret, L. & Amar, M. ben. The radial growth phase of malignant melanoma: multi-phase modelling, numerical simulations and linear stability analysis. J R Soc Interface 8, 345 (2011).

44. Zhang, H., Cisse, M., Dauphin, Y. N. & Lopez-Paz, D. mixup: Beyond Empirical Risk Minimization. Preprint at https://github.com/facebookresearch/mixup-cifar10. (2022).

45. Brunner, A. et al. Ultra-high sensitivity mass spectrometry quantifies single-cell proteome changes upon perturbation. Mol Syst Biol 18, (2022).

46. Derks, J. et al. Increasing the throughput of sensitive proteomics by plexDIA. Nature Biotechnology 2022 1–10 (2022) doi:10.1038/s41587-022-01389-w.

47. Huffman, R. G. et al. Prioritized single-cell proteomics reveals molecular and functional polarization across primary macrophages. bioRxiv 2022.03.16.484655 (2022) doi:10.1101/2022.03.16.484655.

48. Leduc, A., Huffman, R. G. & Slavov, N. Droplet sample preparation for single-cell proteomics applied to the cell cycle. doi:10.1101/2021.04.24.441211.

49. Dou, M. et al. High-Throughput Single Cell Proteomics Enabled by Multiplex Isobaric Labeling in a Nanodroplet Sample Preparation Platform. Anal Chem 91, 13119–13127 (2019).

50. Beck, L. et al. Clinical Proteomics of Metastatic Melanoma Reveals Profiles of Organ Specificity and Treatment Resistance. doi:10.1158/1078-0432.CCR-20-3752.

51. Wolf, F. A., Angerer, P. & Theis, F. J. SCANPY: Large-scale single-cell gene expression data analysis. Genome Biol 19, 1–5 (2018).

52. Sherman, B. T. et al. DAVID: a web server for functional enrichment analysis and functional annotation of gene lists (2021 update). Nucleic Acids Res 50, W216–W221 (2022).

